# Kinetic Modeling of West Nile Virus Fusion Indicates an Off-pathway State

**DOI:** 10.1101/2020.06.05.132605

**Authors:** Abraham Park, Robert J. Rawle

## Abstract

West Nile virus (WNV) is a prominent mosquito-borne flavivirus that causes febrile illness in humans. To infect host cells, WNV virions first bind to plasma membrane receptors, then initiate membrane fusion following endocytosis. The viral transmembrane E protein, triggered by endosomal pH, catalyzes fusion while undergoing a dimer-to-trimer transition. Previously, single-particle WNV fusion data was interrogated with a stochastic cellular automaton simulation, which modeled the E proteins during the fusion process. The results supported a linear fusion mechanism, with E protein trimerization being rate-limiting. Here, we present corrections to the previous simulation, and apply them to the WNV fusion data. We observe that a linear mechanism is no longer sufficient to fit the data. Instead, an off-pathway state is necessary; these results are corroborated by chemical kinetics modeling. When compared with a similar Zika virus fusion model, this suggests that off-pathway fusion mechanisms may characterize flaviviruses more broadly.

## Introduction

Flaviviruses, including West Nile virus (WNV), Zika virus, and dengue virus, are a family of icosahedral, membrane-enveloped, positive sense single-stranded RNA viruses that are responsible for substantial morbidity and mortality worldwide (Barrows et al., 2018; Chong et al., 2019; Diamond, 2009; Heinz and Stiasny, 2012). As with many other membrane-enveloped viruses, the initial steps of infection for flaviviruses are 1) attachment or binding to the host cell plasma membrane and 2) subsequent membrane fusion of the viral envelope with a host membrane at varying stages of endocytosis, depending on the virus (Kaufmann and Rossmann, 2011; Krishnan et al., 2007; van der Schaar et al., 2007). The viral transmembrane E protein, a class II fusion protein existing as closely-packed dimers on the surface of mature virions, is involved in both these steps, and it is essential for facilitating membrane fusion (Harrison, 2015; Perera-Lecoin et al., 2013). The prevailing model (Harrison, 2015; Kaufmann and Rossmann, 2011) for flavivirus fusion suggests that upon exposure to low pH during endocytosis, E protein monomers disassociate from their dimer partner, rearrange to an extended conformation and insert their hydrophobic fusion loop into the host membrane, and then trimerize with nearby extended monomers. Multiple trimers acting together can mediate fusion between the viral and host membranes, refolding into a postfusion conformation with hydrophobic fusion loops brought into close proximity with transmembrane domains. Various sets of structural, kinetic, and simulation data have explored different aspects of this model (reviewed in (Kaufmann and Rossmann, 2011)), yet many of the features (especially of the intermediate transitions) remain murky due to the complex nature of the fusion process which involves many lipid and protein players acting across multiple relevant timescales.

As with other virus families (Floyd et al., 2008; Ivanovic et al., 2013; Kim et al., 2017; Liu and Boxer, 2020; Otterstrom et al., 2014; Rawle et al., 2016; Yang et al., 2015, 2017), single virus fusion experiments have been applied fruitfully to study fusion of flaviviruses (Chao et al., 2014, 2018; Rawle et al., 2018). In these studies, single virus fusion events are collected by fluorescence microscopy, typically observing pH-triggered lipid exchange between fluorescently labeled virions or virus-like particles (VLPs) and model lipid membranes as surrogates for the host cell membrane. The distributions of individual event waiting times between pH drop and the onset of lipid mixing are then analyzed by kinetic modeling. Such modeling efforts have previously been applied to a variety of viruses to estimate the numbers of fusion proteins involved in the fusion process, to identify or enumerate rate limiting steps, and to suggest dependence of the rate-limiting steps on viral structural features which can then be probed by mutagenesis (Chao et al., 2014, 2018; Floyd et al., 2008; Ivanovic and Harrison, 2015; Ivanovic et al., 2013; Rawle et al., 2018).

In this report, we present an advancement to previous kinetic modeling of WNV single virus fusion data by Chao et al.). In the previous modeling approach, the authors used a stochastic cellular automaton (CA) simulation to model their fusion data, and concluded that fusion could be modeled as a linear process with E protein trimerization being the rate-limiting step. In adapting that simulation to model our own Zika virus fusion data in a separate report (Rawle et al., 2018), we discovered shortcomings with the simulation model (detailed in the Materials and Methods). After addressing those shortcomings in a new implementation of that simulation, we discovered that the Zika virus fusion data could only be fit with models containing an additional off-pathway state. However, the updated cellular automaton simulation was not the primary focus of our study. Here, we more directly present the updates to the Chao et al. simulation model, and then apply it to fit their WNV fusion kinetic data, to which they have generously given us access. Strikingly, we observe that when the updated simulation is used to fit the WNV kinetic data, an off-pathway model is needed to successfully fit the data, similar to our previous observations with Zika virus. This opens the possibility that an off-pathway state may be a feature of the flavivirus fusion mechanism more broadly.

## Results and Discussion

### Description of the experimental data

A detailed description of the single virus fusion experimental system and data used for kinetic modeling is reported in Chao et al (Chao et al., 2014). Briefly, DiD-labeled WNV VLPs were bound to glass-supported bilayers using membrane-anchored pseudo-receptors inside a microfluidic device. The membrane-bound pseudo-receptors were either the E16 Fab, which binds to domain III of the WNV E protein, or the lectin domain of DC-SIGN-R. After 5 minutes of VLP binding, fusion was triggered by infusion of a low pH buffer mimicking endosomal pH. Hemifusion was observed by total internal reflectance fluorescence microscopy as lipid exchange between the fluorescently-labeled VLP envelope and the supported bilayer. The waiting time between pH drop and hemifusion for individual VLPs was quantified via image processing. The waiting times from many particles were then compiled into histograms or cumulative distribution functions. Efficiency (also called final extent) was calculated as the number of observed hemifusion events over the total number of particles observed. Their data is replotted in many of the results figures throughout this manuscript (e.g. open circles and bars in Figures 2, 3, S1). We note that cumulative distribution functions are preferred over histogram presentations so as to avoid bias in the data presentation through the choice of bin width, as has been reported (Rawle et al., 2018).

### Description of the cellular automaton simulation algorithm

The cellular automaton simulation models single virus fusion events by modeling the E proteins at the virus-target membrane interface as a close-packed hexagonal lattice (Figure 1). The lattice consisted of 30 monomers, corresponding to ~½ of the E monomers on a typical WNV VLP as described in (Chao et al., 2014). E proteins start in their natively folded state (State F), in which they exist as pre-arranged dimers. Each monomer can individually be activated to an extended state (State E) with pH-dependent kinetics, modeling fusion loop insertion into the target membrane. Dimer partners can be treated with positive cooperativity; if one partner adopts an extended state, the other partner is destabilized and is more likely to be activated. Extended E protein monomers can trimerize (State T) with two neighboring activated monomers, also in a pH-dependent process. If a minimum number of adjacent trimers exist (N_tri_, found to be 2 by Chao et al.), then the virus can undergo hemifusion (lipid mixing, State HF). The waiting time between the start of the simulation (defined as the time of pH drop) and hemifusion is recorded. This process is repeated for hundreds of individual viral particles at the experimentally tested pH values, and the recorded waiting times are compiled into a cumulative distribution function, modeling single virus fusion event data. Finally, for simplification in the rest of the manuscript, we represent the connectivity between states in the simulation by a kinetic model diagram as shown in Figure 1D, but we note this does not capture the geometrical interactions between monomers which are required for certain states to be accessible.

**Figure 1.**
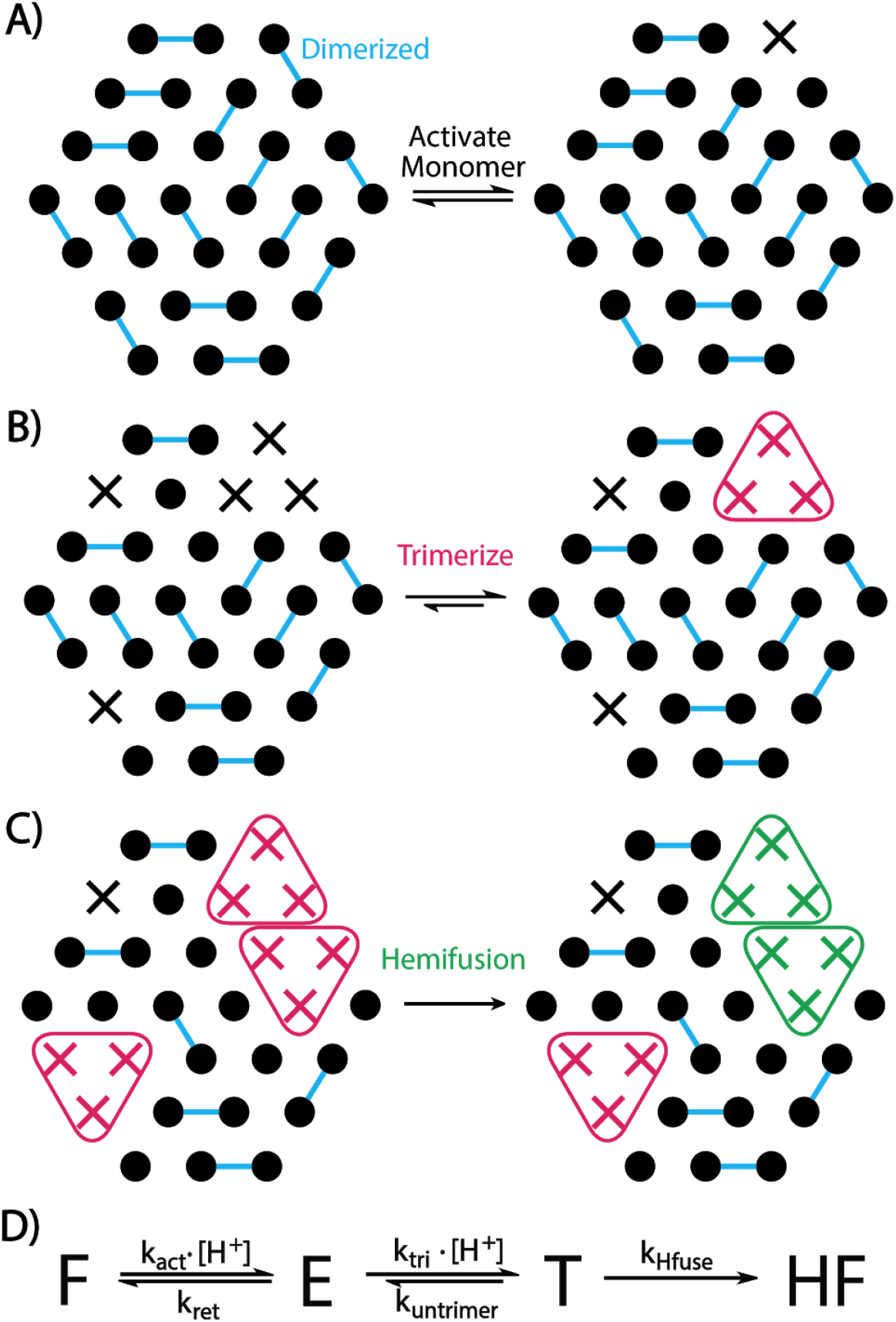
Schematic of cellular automaton simulation of WNV hemifusion. The viral E proteins for a single virus-target interface were modeled as a hexagonal array. At the start of the simulation, each protein monomer started in the Folded state (black dots) and was defined to have a dimer pair (blue lines show connectivity), mimicking WNV structural models. During the course of the simulation, each protein could transition between various states, depending on the state of its neighbors; examples are shown in panels A-C. A) pH activation causes an E monomer in the folded state to transition to the Extended state (black X), representing fusion loop insertion into the target membrane. This activation process was modeled cooperatively; monomers whose dimer partner was no longer in the folded state became more likely to be activated themselves in subsequent time steps. This was implemented as CF*k_act_, where CF is the cooperativity factor. B) Three adjacent monomers in the extended state can form a Trimer (red Xs and triangle) in a pH-dependent process. C) If a minimum number of adjacent trimers exist (depicted here as 2), they can access the HemiFusion state (green Xs and triangle). Once a modeled virion reaches the hemifusion state, the simulation for that particular virion stops, and the time to hemifusion is recorded. The data from many modeled virions is compiled to compare to experimental single virus fusion data. Panel D shows a simplified kinetic model diagram of the above schematic as used in subsequent figures; this diagram shows the connectivity between states, but does not show the geometrical interactions between monomers that are required for certain states to be accessible. State definitions are: F = folded, E = extended, T = trimer, HF = hemifusion. Generic rate constants are as shown.

A variety of corrections and updates were implemented to the previous cellular automaton simulation reported in Chao et al (Chao et al., 2014), largely dealing with the implementation of the algorithm in the code, not to the algorithm itself as described above. These are detailed in the Materials and Methods section. These corrections and updates had a large influence on the outcome of the simulation. When we ran our updated simulation with the best fit parameters reported by Chao et al (Chao et al., 2014), we observed no fusion events at any pH value tested; all E protein monomers largely remained in the folded state (F) over the time window of the experiment (Figure S1).

### A linear model is not sufficient to fit the WNV data

Using the updated simulation, we asked whether a linear model with new best fit parameters could be used to successfully fit the WNV fusion data. To narrow the parameter search to physiologically relevant values, we first used various data sets to generate estimates of a subset of the parameters which were then kept constant in the final fit, similar to the process used for the Zika virus fusion model (Rawle et al., 2018).

#### Estimating k_Hfuse_

From the pH-dependent WNV fusion data (see Figure S2B, open circles), we see minor differences in the curve shapes between high (pH 5.75-6.25) and low (pH 5.-5.5) pH values, but at low pH values (pH 5-5.5) the curve shapes appear to reach a limiting value. This suggests that the rate limiting step at low pH values is not pH-dependent. In the scheme as drawn, this would be the final hemifusion step (k_Hfuse_). Therefore, we estimated k_Hfuse_ at N_tri_ = 1, 2, or 3 by fitting the pH 5 fusion data to a model where all previous forward steps were given very large rate constants. The best fit estimates for k_Hfuse_ were determined to be 0.0047 s^−1^ for N_tri_ = 1 and 2, and 0.009 s^−1^ for N_tri_ = 3 (Figure S2). We noted that N_tri_ = 3 had a poorer quality fit than N_tri_ = 1 or 2, likely due to the inability of some viral particles to achieve hemifusion because the stochastic pattern of trimers formed in the hexagonal lattice did not allow the minimum number of adjacent trimers (N_tri_ = 3) to be achieved.

#### Estimating k_ret_/k_act_

We used dynamic light scattering data from Chao et al. (Chao et al., 2014) to estimate the ratio of k_ret_ to k_act_. In their report, they measured the equilibrium hydrodynamic radius of Kunjin virus in the absence of target membranes as a function of pH, observing a pKa of 6.8. In the context of the cellular automaton model, this pKa represents the equilibrium between the folded E proteins and extended monomers. Consequently, 10^−6.8^ = k_ret_/k_act_ and k_ret_ = k_act_ * 10^6.8^.

#### Estimatingk_untrimer_

We set k_untrimer_ = 10^−9^ s^-1^, as was done by Chao et al. This is consistent with previous liposome co-flotation data demonstrating that flavivirus trimer formation and fusion loop capture in target liposomes is essentially irreversible (Allison et al., 1995; Chao et al., 2014; Stiasny and Heinz, 2004).

Using these estimated parameters, we then fit the WNV fusion data using the model in Figure 2A, varying k_act_ and k_tri_ with N_tri_ = 1, 2, or 3. Similar to the Zika virus data reported previously (Rawle et al., 2018), we observed that the best fit from the linear model could roughly capture the general trend in final extent, but only at the expense of a very poor fit to the curve shapes (Figures 2C-E, S3). Additionally, allowing the cooperativity factor CF (previously set at 1) to vary in the fit for N_tri_ = 2 did not substantially improve its ability to fit the data (Figure S4). Therefore, we concluded that, as with the Zika virus fusion data, a linear model could not successfully fit the WNV fusion data.

**Figure 2.**
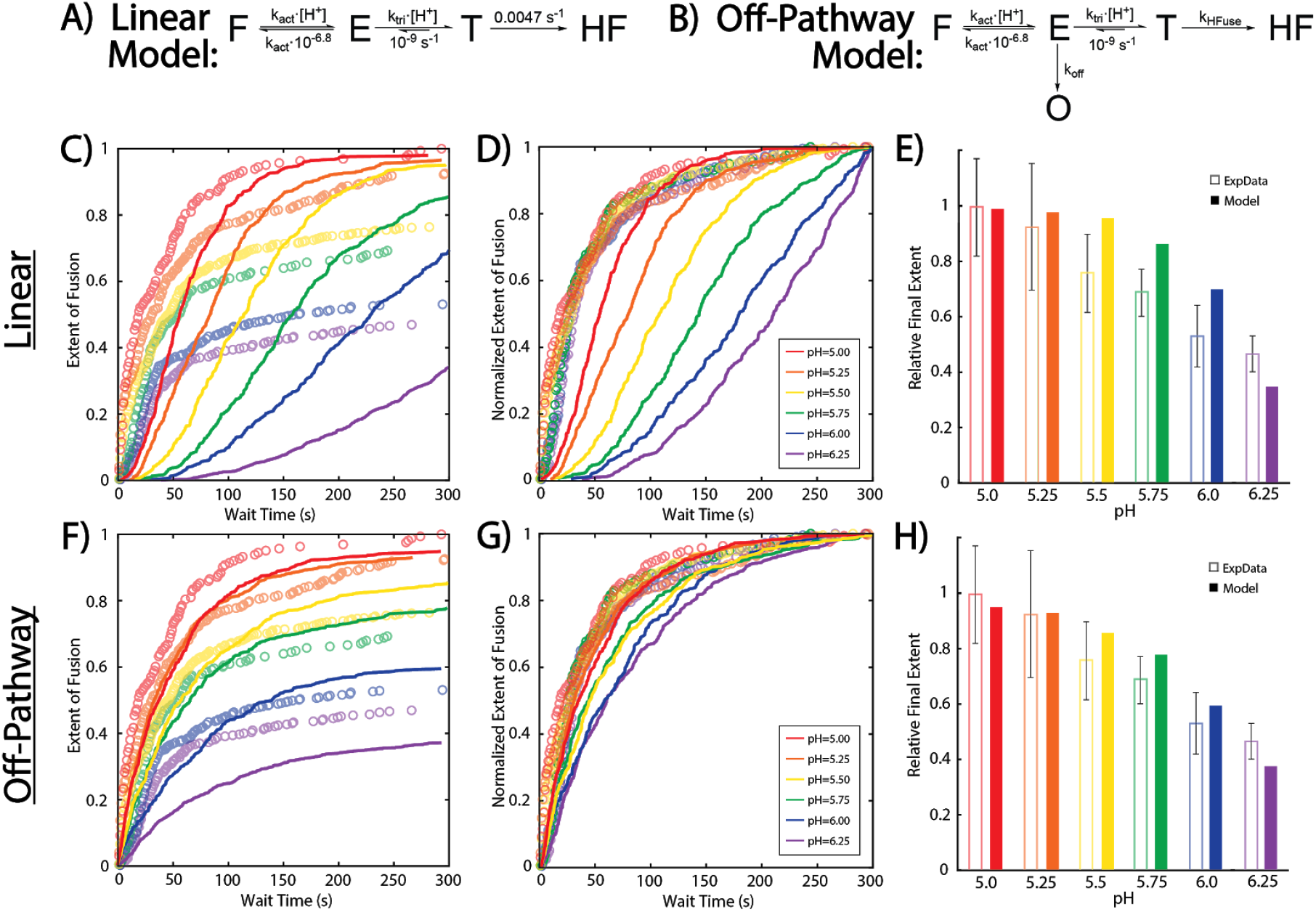
Cellular automaton modeling of WNV kinetic fusion data indicates presence of off-pathway state. Single virus hemifusion data from (Chao et al., 2014) was fit to either a 4-state linear model (A) or a 5-state off-pathway model (B) of the cellular automaton simulation. CDFs of all pH values were fit simultaneously and some fit parameters were constrained to physiologically relevant values as described in the main text. Panels C and F show best fits (solid lines) to the experimental relative extent of hemifusion data (open circles) vs wait time for the linear and off-pathway models respectively. Extents of hemifusion were calculated relative to the final extent at pH 5, which was set at 1. Panels D and G show the normalized fits overlaid on the normalized experimental data to demonstrate how well the fits model the experimental curve shapes. Simulation and experimental data were normalized to the final extent of hemifusion at each pH value. Panels E and H show the best fit model predictions at each pH value for final extents of fusion (solid bars) at t = 300 sec compared to the experimental final extents (open bars). Final extents are shown relative to pH 5, which was set 1. Error bars are as reported by Chao et al., standard deviation of 3-5 independent experiments. In all cases, the best fit off-pathway model showed an improved fit to the experimental data as compared to the linear model. Best fit parameters for the linear model shown are: k_act_ = 10^5^ M^−1^ s^−1^, k_tri_ = 1000 M^−1^ s^−1^, N_tri_ = 2, and CF = 1; and for the off-pathway model are: k_act_ = 10^8^ M^−1^s^−1^, k_tri_= 10^6^ M^−1^s^−1^, k_off_= 1.2 s^−1^, k_Hfuse_= 0.0047 s^−1^, N_tri_= 2, and CF = 1.

### An off-pathway state model is sufficient to fit the WNV data

We then examined whether the addition of an off-pathway state would enable the simulation to successfully fit the WNV data. We used a similar rationale as in the development of our previous model for Zika virus fusion (Rawle et al., 2018); the branching between the off-pathway state and the forward reaction will set an upper limit to the final efficiency (extent) of the reaction. If the immediate forward reaction step is pH-dependent, this will ensure that the final efficiency is pH-dependent, as is observed in the data. The normalized rates (or curve shapes), on the other hand, will be largely determined by the rate-limiting steps in the forward reaction.

We positioned the off-pathway state as branching from the extended monomer state (Figure 2B). This position is not necessarily required; it is likely that the off-pathway state could be branched off of an additional pH-dependent state in between the F and E states with a similar outcome. However, to construct the most parsimonious model, we only chose to add 1 additional state.

With this off-pathway model, we determined the best fit to the WNV data. We used identical physiologically relevant estimates for a subset of parameters as in the linear model (e.g. k_Hfuse_ = 0.0047 s^−1^, 0.0047 s^−1^ or 0.009 s^−1^; k_ret_ = k_act_ * 10^−6.8^; N_tri_ = 1, 2 or 3; k_untrimer_ = 10^−9^ s^−1^). As a limiting case, we also treated the off-pathway state as irreversible. The return from the off-pathway state must necessarily be slow relative to k_off_, otherwise the model becomes nearly identical to the linear model, with very little time spent in the off-pathway state (Figure S5). k_tri_, k_act_, and k_off_ were varied independently with N_tri_ = 1, 2 or 3. Using this off-pathway model (Figure 2B), we observed an improved fit to the WNV data (Figure 2 F-H). Curve shapes were largely maintained across the pH range, and the simulated efficiencies fell within error estimates for all pH values except 5.75 and 6.25. This suggests that, as with previous Zika virus fusion data, an off-pathway state is needed to successfully model WNV fusion data.

We also observed that N_tri_ = 2 showed a marginal improvement in fit quality over N_tri_ = 1 within the stochastic noise limit of the simulations (Figure S6). For N_tri_ = 3 the curve shapes more closely matched the data (Figure S6, panel H), but the efficiency change with pH was poor, likely due to the inability of a large percentage (~15%) of particles to form the required number of trimers (see also Figure S2), lowering the upper bound on the maximum possible efficiency (Figure S6, panel G). This suggests that given the assumptions of the simulation model, the minimum number of trimers needed to mediate hemifusion is less than 3.

Additionally, we observed that the fit was relatively insensitive to the value of k_act_, as long as it was much larger than k_tri_. Within the model, this indicates that the rate-limiting step is either trimerization or hemifusion, depending on the pH value. Furthermore, this implies an insensitivity to the cooperativity factor CF, which only affects the F -> E transition. Indeed, when we varied CF in addition to k_tri_, k_act_, and k_off_ in the fit at N_tri_ = 2, there was little improvement for values of CF > 1 (Figure S7).

We also experimented with making the return from the off pathway reversible in a more limited fashion (Figure S5). In the context of the best fit parameters for N_tri_ = 2, we varied the rate constant of return from the off-pathway state (k_return_). We observed that values of k_return_ < 0.01 * k_off_ were identical within the stochastic noise limit, but larger values approached the qualitative behavior for the linear model. This supports the assertion that k_return_ must necessarily be slow relative to k_off_.

### Per virus kinetic modeling also indicates that an off-pathway state is needed

We also examined the ability of a model inspired by chemical kinetics, either with or without an off-pathway step, to successfully fit the WNV data, as had been done for the Zika virus data (Rawle et al., 2018) (Figure 3). As compared with the cellular automaton, which models individual E proteins at the virus-target membrane interface, this chemical kinetics-inspired approach models the entire virion as it transitions between generalized states. In the most parsimonious linear version of this model, each virion can exist in 3 states: B) bound to the target membrane prior to pH activation, A) pH-activated, and HF) hemi-fused with the target membrane. All virions start at the B state at t = 0, which is the time at which pH drop occurs. Similar to the development of the cellular automaton model, we first estimated some parameters at physiologically relevant values. k_AHF_. was estimated by fitting the pH 5 hemifusion data to a single exponential (best fit k_AHF_ = 0.024 s^−1^), and k_AB_ was estimated as k_BA_*10^−6.8^ using the Kunjin virus hydrodynamic radius data from Chao et al. (Chao et al., 2014).

**Figure 3.**
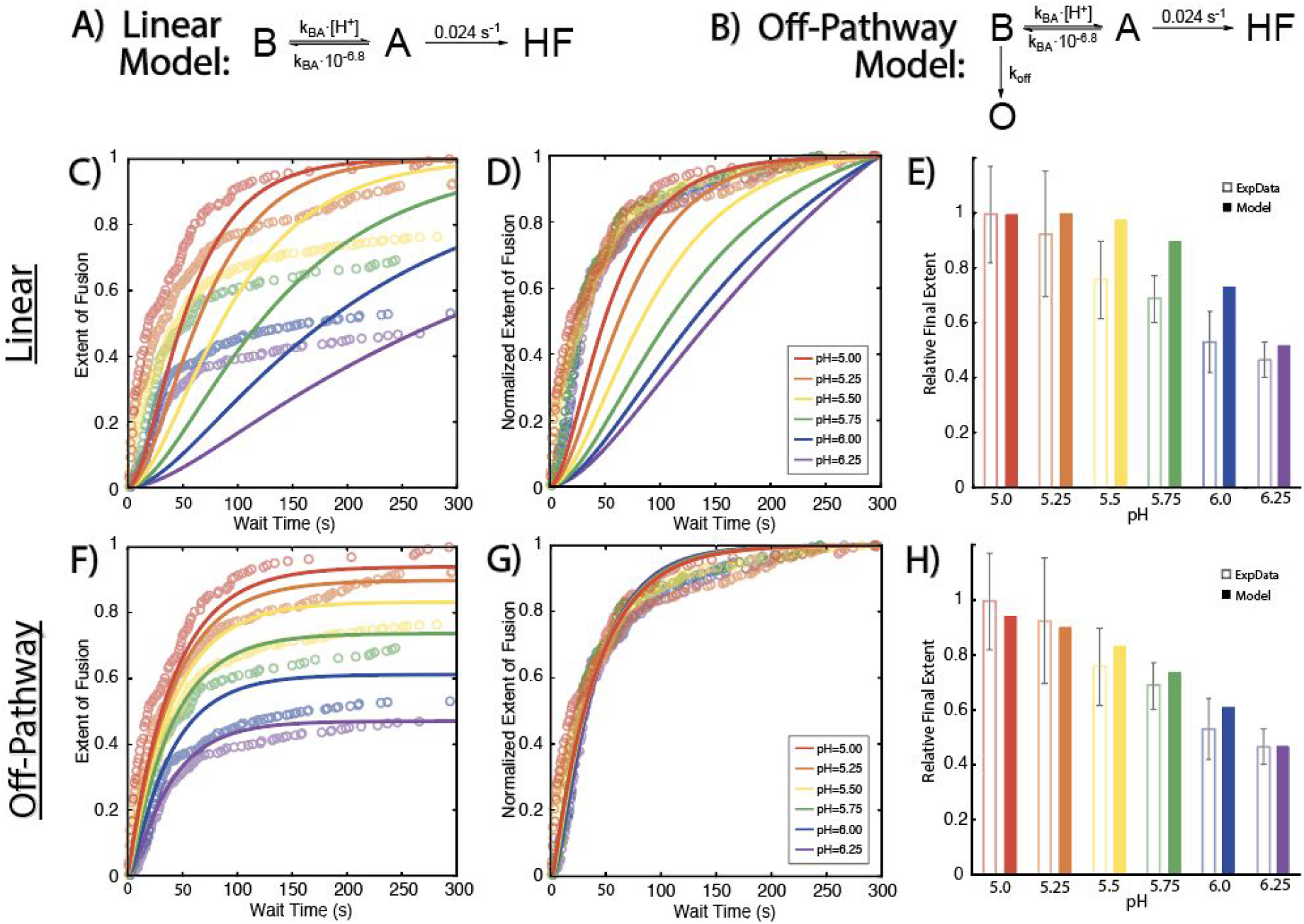
Per virus kinetic modeling also indicates presence of off-pathway state. Single virus hemifusion data from (Chao et al., 2014) was fit to either a 3-state linear (A) or a 4-state off-pathway (B) model, using chemical kinetics-inspired modeling of the entire virion. CDFs of all pH values were fit simultaneously and some fit parameters were constrained to physiologically relevant values or relative to other rate constants as described in the main text. Panels C and F show best fits (solid lines) to the experimental relative extent of hemifusion data (open circles) vs wait time for the linear and off-pathway models respectively. Extents of hemifusion were calculated relative to the final extent at pH 5, which was set at 1. Panels D and G show the normalized fits overlaid on the normalized experimental data to demonstrate how well the fits model the experimental curve shapes. Model best fits and experimental data were normalized to the final extent of hemifusion at each pH value. Panels E and H show the best fit model predictions at each pH value for final extents of fusion (solid bars) at t = 300 sec compared to the experimental final extents (open bars). Final extents are shown relative to pH 5, which was set 1. Error bars are as reported by Chao et al.. In all cases, the best fit off-pathway model showed an improved fit to the experimental data as compared to the linear model. The best-fitting rate constants for the linear model were: k_BA_ = 5330 M^−1^s^−1^; and for the off-pathway model were: k_BA_=1.5 x 10^5^ M^−1^s^−1^ and k_off_= 0.05 s^−1^.

With these estimates, the WNV data was then fit to the linear model (Figure 3A), varying k_BA_. As with the cellular automaton simulation, we observed that the linear model could not successfully fit the WNV data (Figure 3C-E). Indeed, we propose that any linear model cannot successfully fit the WNV data, even if additional steps are added. If additional steps are added which are not rate-limiting, they will not appreciably alter the fit as shown. If additional rate-limiting steps are added, then a large lag phase will be introduced, which does not match the observed kinetics (Rawle et al., 2018). Furthermore, the only way in which a linear model can successfully capture a pH-dependent decrease in efficiency is for the extent of fusion to not level off into a plateau phase as shown in Figure 3C-D (solid lines); the experimental data (open circles) however clearly shows the extent leveling off at all pH values. Similar results were observed previously in the modeling of Zika virus data (Rawle et al., 2018).

We then added an off-pathway state O to the model, branching off the B state. As with the off-pathway cellular automaton model, the branching between the pH-dependent BA transition and the pH-independent BO transition enforces an upper bound to the efficiency which can be achieved. Meanwhile, the forward rate constants govern the curve shapes. Again, all virions start in the B state at t = 0. As with the CA model, this assumes that the off-pathway state is only accessible following target membrane binding and pH activation.

Using this off-pathway model (Figure 3B), we observed a high-quality fit to the WNV data (Figure 3F-H). Curve shapes remained largely invariant across the pH range, and efficiencies fell within error estimates of the data for all pH values. This further supports the conclusion from the cellular automaton simulation results - an off-pathway state is needed to successfully fit the WNV data.

Additionally, we also tested the influence of reversibility of the off-pathway state on the model (Figure S8). We observed that the best fit for the reversible off-pathway model was largely similar to the irreversible model, with some improvement to fitting the curve shapes. For the reversible model, the best fit for k_return_ was ~500X lower than k_off_, consistent with the observation from the CA simulation results (Figure S5) that k_return_ must be much smaller than k_off_ or else the model approaches the linear model, with little time spent in the off-pathway state.

We also note, analogous to the cellular automaton model, that the above presentation is for the most parsimonious off-pathway model, only having 4 states. Additional linear states may be added to the model, but they must necessarily be fast relative to the states as described above otherwise a lag phase is introduced.

### Limitation for both models

Finally, we note that both sets of off-pathway models (cellular automaton and per virus) suffer from a similar limitation. As implemented herein, both models define the start state of the system at t = 0 to be either in the Folded state (cellular automaton) or State B (per virus modeling). If however the models are extended to simulate the time between virus binding to the target membrane and pH drop, during which time the virus is bound to the target membrane at pH 7.4, then the models predict a very strong dependence on the length of that time window. In essence, the models predict a significant fraction of viral proteins/virions end up in the off-pathway state prior to pH drop, which limits the possible final extent. In the WNV experimental data, virions were bound during 5 minutes and this time was not varied. Consequently, there is no data to indicate the sensitivity of that binding time window on the observed efficiency, and therefore it was not modeled here. However, in the Zika virus data, this time window was varied, with only minor sensitivity to the binding time window being observed. If we assume that the same holds true for the WNV data, this suggests that both sets of models are not capturing an important physical parameter.

## Conclusion

The cellular automaton and chemical kinetics modeling results indicate that an off-pathway state is necessary to successfully fit WNV hemifusion data. Taken together with the previous Zika virus data, which supported a similar conclusion, these results raise the possibility that an off-pathway state may be a feature of the flavivirus fusion mechanism more broadly.

As of this report, there is insufficient biochemical and structural data to definitively identify this off-pathway state. However, inferences from the single particle fusion data and the kinetic modeling can provide constraints and hypotheses. First, the off-pathway state must branch prior to a pH-dependent step, otherwise no pH-dependence in the final extent would be observed. This suggests that the off-pathway state occurs prior to trimerization. Second, the models we employ suggest that the off-pathway state observed here may be dependent on target membrane insertion. This suggests that it may be distinct, at least in some respects, from reversible exposure of cryptic epitopes (“viral breathing”) or from slow viral inactivation in solution, also termed intrinsic decay, which has been reported for several flaviviruses (Dowd et al., 2014; Kuhn et al., 2015). We cannot, however, rule out that the structural features of the off-pathway state reported here may be related. Finally, the two observations above raise the possibility that this off-pathway state occurs following pH-activation of E monomers; perhaps it may be a misfolded or aggregated state of the E protein following membrane insertion. This hypothesis would suggest that the off-pathway state may be distinct from the states stabilized by small molecule flavivirus E protein inhibitors, which have been proposed to inhibit fusion loop exposure (Chao et al., 2018; de Wispelaere et al., 2018) and/or stabilize trimers in a fusion incompetent state (Schmidt et al., 2012). However, the modeling results shown here do not definitively rule out those possibilities.

Finally, we also make a high level observation about the robustness of the experimental data used as the basis for kinetic modeling (Chao et al., 2014; Rawle et al., 2018). Both the Zika and WNV data sets in broad strokes show similar features – hemifusion occurs on similar timescales, curve shapes are largely maintained across the pH values tested, and a roughly linear decrease in efficiency with increasing pH is observed. Interestingly, such similarities in the data are observed in spite of key differences in the experimental setups. 1) The Zika data was collected with infectious virus, whereas the WNV data used VLPs. 2) The Zika data used tethered liposomes as target membranes for fusion, whereas the WNV data used planar supported lipid bilayers. 3) Different surrogate receptors were used: DNA-lipid conjugates in the Zika data and membrane-anchored Fabs in the WNV data. 4) Different membrane dyes were used to label the viral envelope: Texas-Red DHPE (Zika) or DiD (WNV). Each of these factors might not unreasonably be expected to bias the observed hemifusion rates or efficiencies (see for example (Rawle et al., 2019)). Yet similar data is observed and similar off-pathway kinetic models fit each data set; this speaks to the robustness of the data and further supports the idea that the off-pathway state is a feature of both Zika and WNV hemifusion, and perhaps of flaviviruses more broadly.

## Materials & Methods

### Modeling of WNV Fusion Data

Cellular Automaton simulations and kinetic models previously used to model Zika Virus fusion (Rawle et al., 2018) were updated for use with WNV data. They were written and implemented in MATLAB R2019a (MathWorks), and are available at https://github.com/rawlelab/WNVmodeling. Specifics for each are described below.

### Cellular Automaton (CA) Simulation

In the CA simulation, single virus-target membrane interfaces were modeled as a hexagonal lattice of viral E protein monomers. The lattice consisted of 30 monomers, corresponding to ~½ of the E monomers on a typical WNV VLP as described (Chao et al., 2014). Each monomer had a defined dimer partner at the beginning of the simulation, and could transition between various states in the model only when certain conditions between neighboring monomers were met. Cooperativity of activation between dimer partners was modeled with a cooperativity factor (CF) as CF * k_act_. Three adjacent activated monomers were required to access the trimerized state, and N_tri_ number of adjacent trimers were required for the viral particle to undergo hemi-fusion. For the trimerization step, each of the adjacent monomers could initiate trimerization independently. Each protein started in the natively folded state (State F) at time t = 0 s. Each virus was modeled for 300 s (the time window of the experimental data) or until it reached hemifusion (State HF), whichever came first. State transition probabilities for each monomer were calculated by solving the system of coupled ordinary differential equations (ODEs), constrained by the current states of neighboring monomers or trimers. At each discrete time step, monomers were evaluated in a random order to determine which transitions would occur. For each pH value, simulations were repeated in groups of 500 virions until at least 200 virions fused or until 1,500 virions had been simulated. The time step of the simulation was set such that the maximum transition probability for any transition would be 0.1. The minimum allowed time step was 100 ms. In benchmarking simulations where the minimum allowed time step was lowered to 10 ms, little difference was observed in the output of the simulation.

Some parameters in the model (k_act_, k_ret_, k_Untrimer_, k_Hfuse_) were estimated at physiologically relevant values using experimental data, as described in the Results and Discussion. The remaining parameters were varied independently, and the best fit was determined using a grid-based search algorithm and Negative Log Likelihood (NLL) minimization. Given that the splitting between trimerization and the off-pathway step is what determines the pH-dependent efficiency changes, we scanned k_off_ in log space as k_off_ = k_tri_*10^−pKoff^ where pK_off_ is equivalent to the pH value where the forward rate of trimerization is equal to the off-pathway rate if all monomers were extended. Ranges scanned for each parameter were: k_tri_ 1,000 to 10,000,000 s^1^, pK_off_ 5.3 to 7.8 s^−1^, N_tri_ 1 to 3, k_act_ 10,000 to 100,000,000 s^−1^, CF 1 to 300. In all cases, the best fitting parameters had converged to within the stochastic noise limit of the simulations. The NLL minimization formulation has been described in detail previously (Rawle et al., 2018).

### Corrections and Updates to the CA Simulation

A variety of corrections and updates were implemented to the previous cellular automaton simulation reported in Chao et al (Chao et al., 2014). These corrections are included in the description of the CA simulation above, but are elaborated below for clarity. The most important correction dealt with how the pH drop was modeled. In the prior simulation, all E protein monomers were instantaneously equilibrated across the folded (F) and extended (E) states at the target pH following pH drop (i.e. at t = 0). For low pH values, this meant that nearly all monomers started in the extended (E) state at t = 0. However, given the values of the rate constants employed (e.g. k_act_ = 1 M^−1^ s^−1^), and the pH values simulated (pH 5 - 6.25), the time to reach equilibration would have been longer than the 300 sec observed experimental time window (e.g. at pH 5, k_ext_ = [H^+^] * k_act_ = 1 x 10^−5^ s^−1^). In the updated simulation therefore, no instantaneous equilibration at the target pH took place following pH drop. Each protein monomer was allowed to transition between states starting from the folded (F) state.

Another update to the simulation improved the calculation of transition probabilities. In the previous simulation, only single transition probabilities were accounted for during a given time step. For instance, a given monomer could transition from the folded (F) to the extended (E) state during a given time step, but it could not also transition back to the F state during the same time step, even if the time step was large relative to the rate constants in question. Consequently, some transitions were undercounted relative to others, which could lead to bias in the simulation outcome. To address this issue in the updated simulation, a matrix exponential formulation was used to solve the system of coupled ordinary differential equations that describe the probabilities of all possible transitions during a given time step. The final step to hemifusion, however, was still only accessible from the trimer state once N_tri_ was achieved; this constraint was necessary to deal with uncertainty as to which neighboring monomers might form a trimer in any given time step.

A smaller correction to the simulation dealt with implementation of the cooperativity factor (CF), which was used to model the destabilizing effect the activation of one monomer could have on its dimer partner. In the previous simulation, the cooperativity factor was implemented by directly multiplying the folded to extended transition probability (i.e. CF * P_fold-ext_). This led to possibilities where the transition probability could be larger than 1 and could lead to an observed insensitivity to the cooperativity factor beyond certain values. In the updated simulation, the CF was implemented by multiplying the k_act_ rate constant, which resolves this issue.

A variety of other smaller corrections were also implemented, such as ensuring that the probabilities of all possible transitions within a single step summed to 1 and that transitions/interactions between monomers were sampled randomly.

Given the variety and scope of these corrections and updates, we kept the central algorithm as outlined in Chao et al (Chao et al., 2014), but rewrote the code implementing the cellular-automaton simulation rather than modifying the pre-existing code. This new implementation also included a variety of updates to improve ease of use for the user. These updates include an algorithm to automatically assign dimer pairs, and a more flexible interface to implement new kinetic models.

### Per Virus Kinetics Modeling

Chemical kinetic modeling was used to model entire virions, rather than individual viral proteins, as they transitioned between different states leading to hemifusion. Modeling was performed by numerically solving the system of coupled ODEs that corresponded to the rate equations of each kinetic model. The solutions were evaluated at discrete time points to determine the number of virions in each state; all viral particles were defined to be in the bound state (State B) at time t = 0 s. For each pH value, the number of viruses that had undergone hemifusion (i.e. reached State HF) at discrete time points was compiled into a cumulative distribution function (CDF).

Some model parameters (k_AHF_, k_AB_) were estimated at physiologically relevant values, as discussed in the Results and Discussion. Initial best guesses for the remaining parameters were estimated by scanning values across several orders of magnitude. We then used an unconstrained Nelder Mead optimization method implemented in MATLAB (Mathworks, Inc) to determine the best fits via NLL minimization. CDFs from all pH values were fit simultaneously. The NLL minimization formulation has been described in detail previously (Rawle et al., 2018).

## Acknowledgments

AP ran simulations and kinetic modeling, did data analysis, and wrote the manuscript. RJR did data analysis, and wrote the manuscript. The authors thank Elizabeth Webster, Katie Liu and Steven Boxer (Stanford University), and Peter Kasson (University of Virginia) for helpful conversations and manuscript feedback. Luke Chao (Harvard University) generously provided access to the WNV experimental data and gave helpful feedback. The authors thank Williams College for financial support.

## Supporting Information

**Figure S1.**
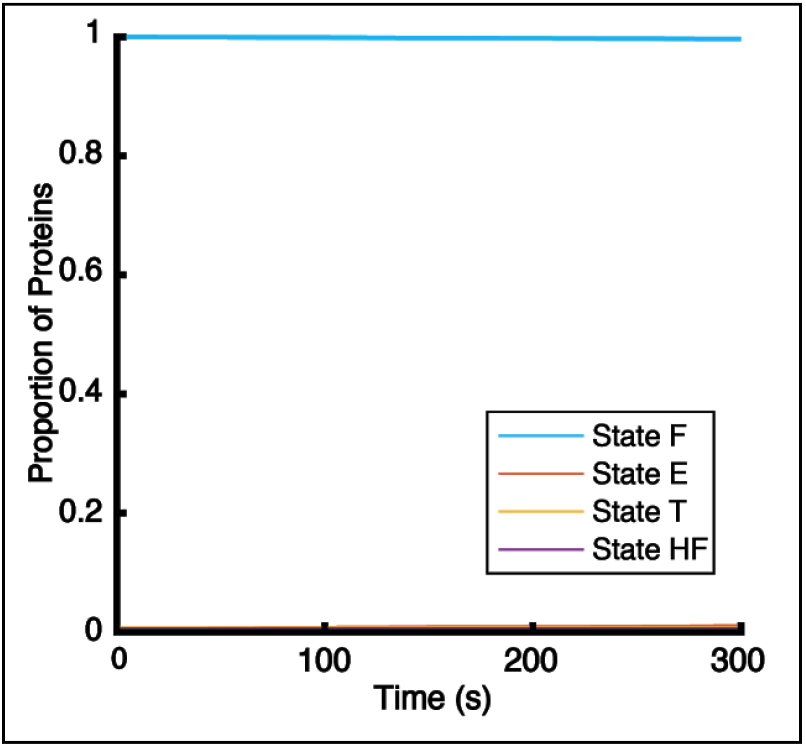
Updated cellular automaton simulation using previous parameters. Best fit parameters from Chao et al. 2014 were used in the updated version of the 4-state linear model cellular automaton simulation. Plotted is the proportion of protein monomers in each state during the 300 sec simulation time window at pH 5, corresponding to the conditions at which the highest efficiency of hemifusion events was observed in the experimental data. All monomers remained in the starting state (State F) for the entirety of the 300 sec simulation; no state transitions or hemifusion events were observed. Parameter values were: k_act_=1 M^−1^s^−1^, k_ret_=1.5 x 10^−7^ s^1^, k_tri_=1 s^−1^, k_Hfuse_=1 s^−1^, N_tri_ = 2, and CF = 1. Number of viral particles simulated was 1,500.

**Figure S2.**
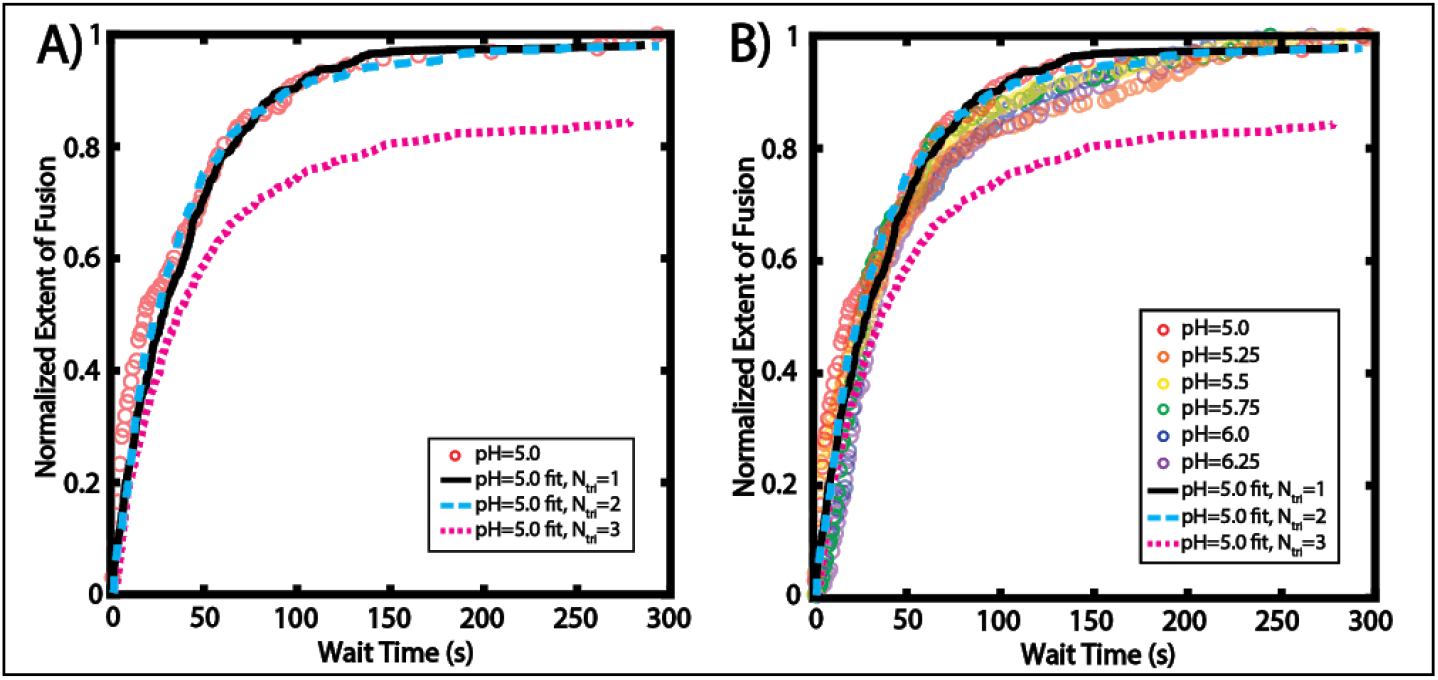
Estimation of k_Hfuse_ using single virus hemifusion data at pH 5. k_Hfuse_ was estimated by fitting the single virus data at pH 5 to the 4-state linear model of the cellular automaton simulation (Fig. 1D). Forward steps prior to the Trimer -> Hemifusion transition were kept very fast (at least 1000x k_Hfuse_), and k_Hfuse_ was independently scanned for each value N_tri_ = 1, 2, or 3. The best-fit rate constants were 0.0047 s^−1^ (N_tri_ = 1), 0.0047 s^−1^ (N_tri_ = 2), and 0.009 s^−1^ (N_tri_ = 3). A) The best fit for N_tri_ = 1, 2, or 3 (solid lines) overlaid on the experimental data CDF at pH 5 (red circles). B) The same best fits overlaid on all the normalized experimental data (open circles) to show minor variations in curve shapes with pH. N_tri_ = 3 exhibited a much poorer fit, likely because some fraction of particles formed a stochastic pattern of trimers that did not allow 3 neighboring trimers to interact, preventing hemi-fusion.

**Figure S3.**
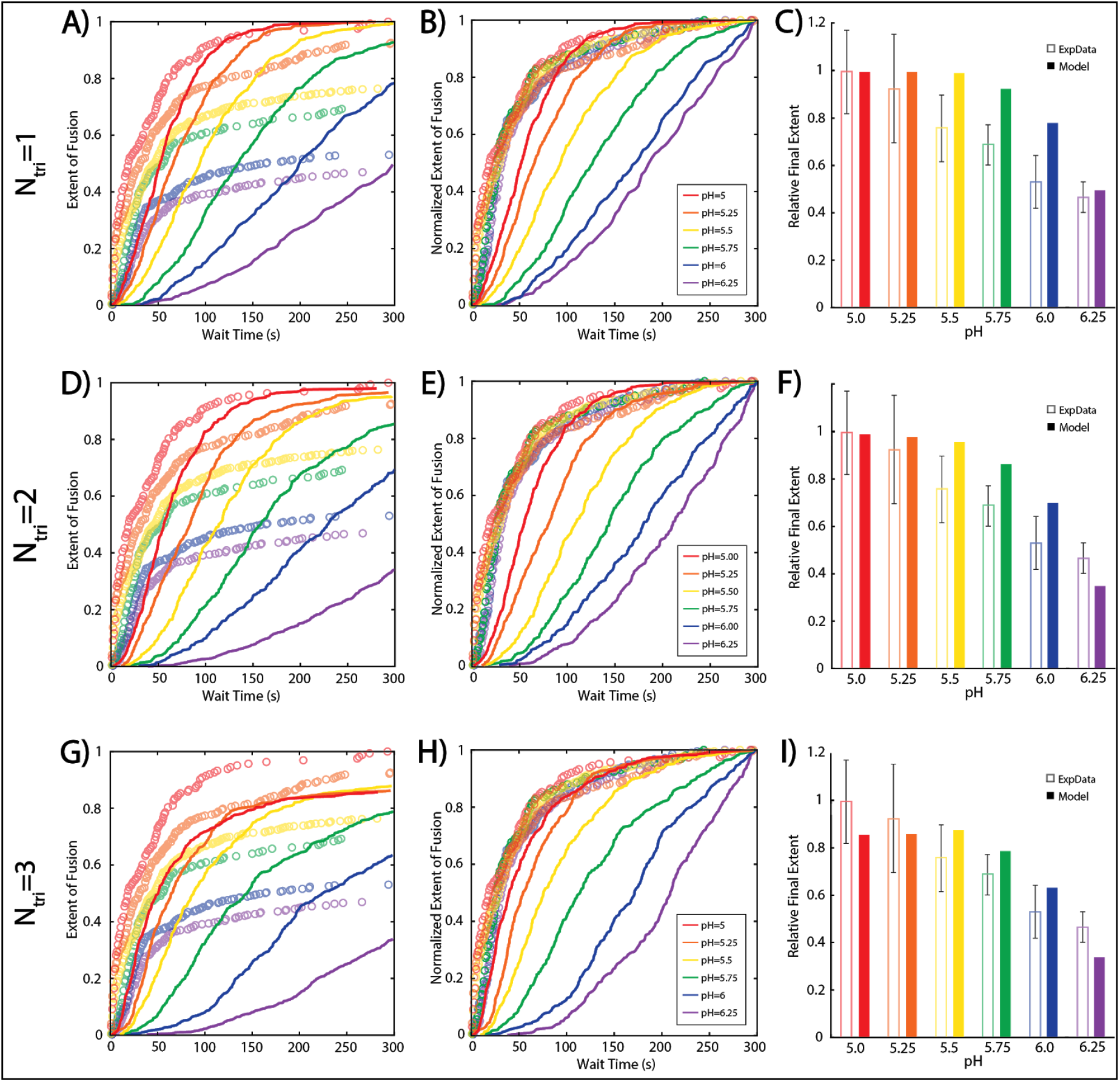
Comparison of CA linear model best fits for N_tri_ = 1, 2, or 3. Single virus hemifusion data was fit to a 4-state linear model of the CA simulation as in Figure 2A. Shown are the best fits for N_tri_ = 1 (Panels A-C) N_tri_ = 2 (Panels D-F), or N_tri_ = 3 (Panels G-I). Best fit parameters for N_tri_ = 1 are: k_act_ = 5 x 10^4^ M^−1^s^−1^, k_tri_= 500 M^−1^s^−1^, k_Hfuse_= 0.0047 s^−1^; for N_tri_ = 2: k_act_ = 10^5^ M^−1^s^−1^, k_tri_= 10^3^ M^−1^s^−1^, k_Hfuse_= 0.0047 s^−1^; for N_tri_ = 3: k_act_ = 3 x 10^5^ M^−1^s^−1^, k_tri_= 3 x 10^3^ M^−1^s^−1^, k_Hfuse_= 0.009 s^−1^.

**Figure S4.**
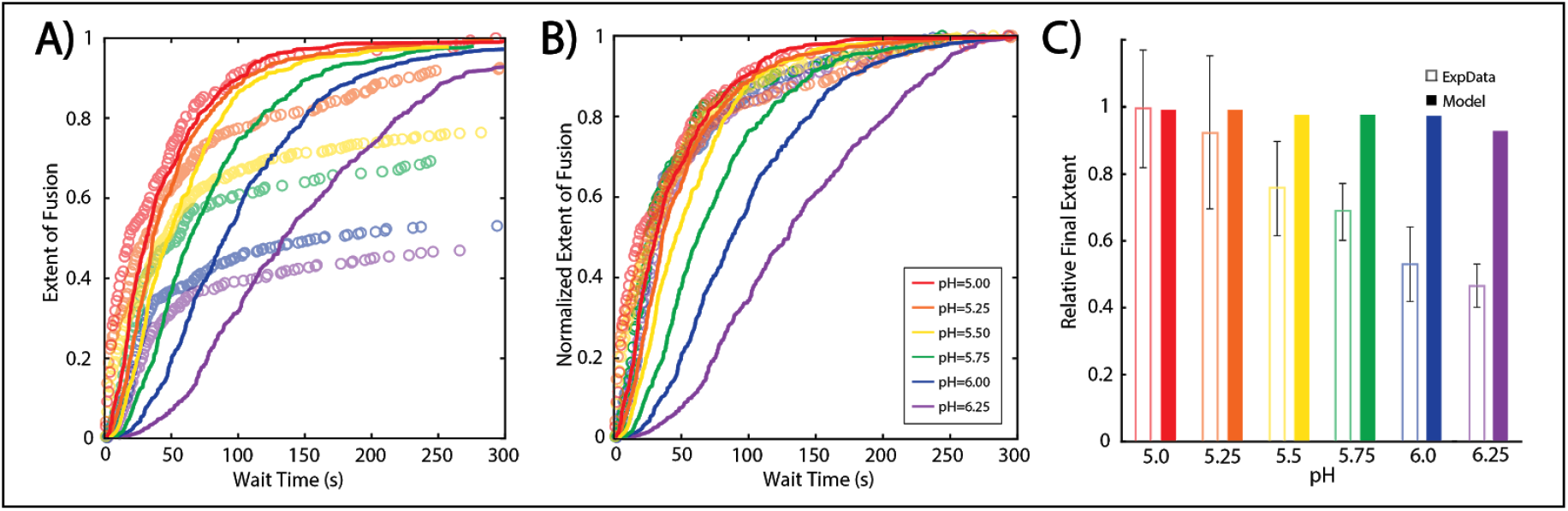
CA linear model best fit at N_tri_ = 2 and varying the Cooperativity Factor. Single virus hemifusion data was fit to a 4-state linear model of the CA simulation as in Figure 2A, but varying the Cooperativity Factor in addition to the other free parameters. Note the marginal improvement in curve shapes (Panel B) relative to the best fit in Figure 2D, but at the expense of very poor fitting to the extent values (compare Panel C and Figure 2E). This is a consistent result with the linear model - it cannot simultaneously match the curve shapes and the trend in final extent. Best-fitting parameters are: k_act_ = 5 x 10^5^ M^−1^s^−1^, k_tri_ = 5 x 10^3^ M^−1^s^−1^, k_Hfuse_ = 0.0047 s^−1^, and CF = 4.

**Figure S5.**
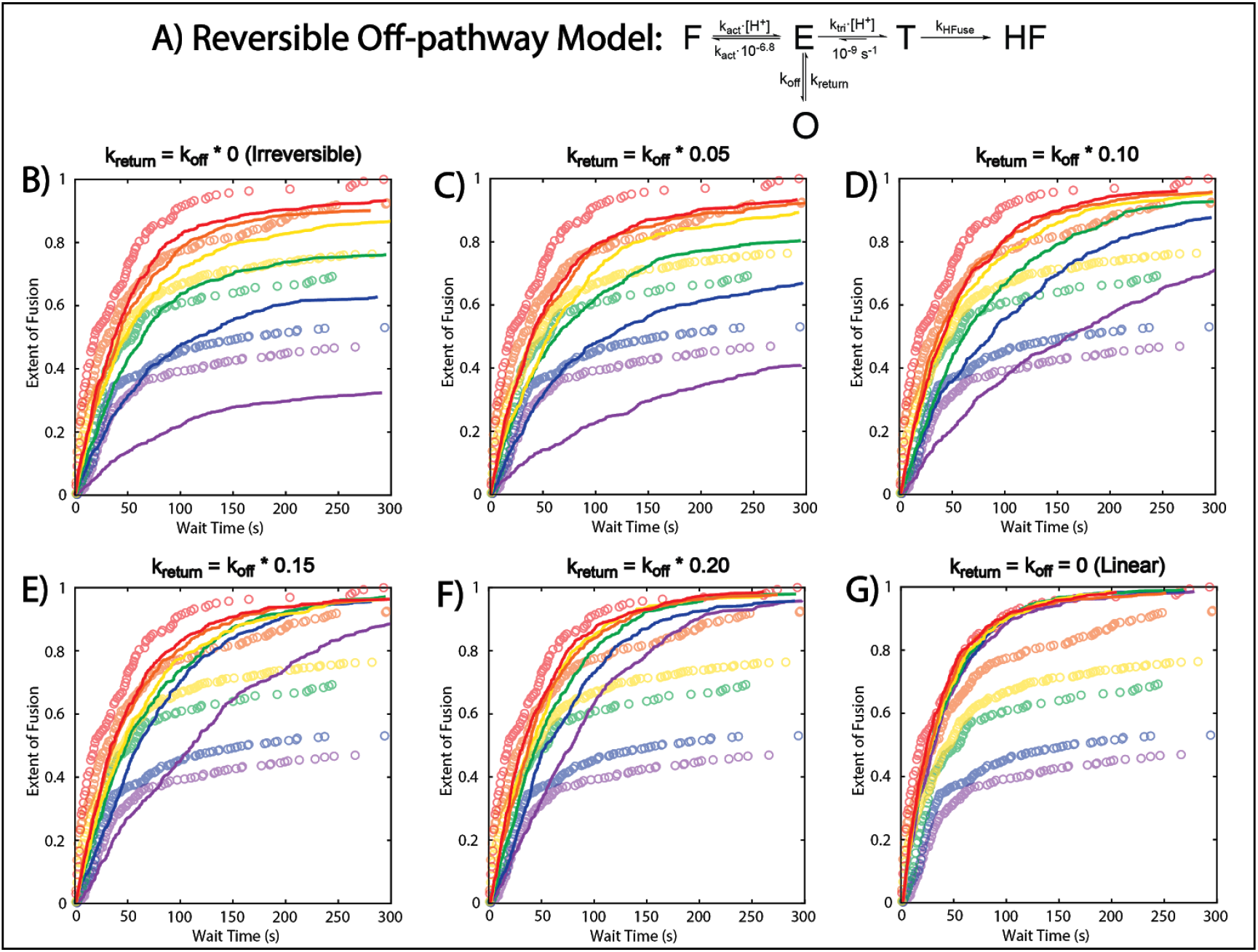
Off-pathway CA-simulated CDFs approach the linear model if k_return_ is similar to k_off_. Reversibility from the off-pathway state was examined by varying k_return_ as a function of k_off_ using the 5-state off-pathway model (Panel A). The results of the simulation (solid lines) are shown over the experimental data (open circles) for selected values of k_return_ (Panels B-F). Panel G shows the simulation results when k_off_ = k_return_ = 0 s^−1^, equivalent to the corresponding linear model. All other parameters used in the graphs displayed are those for the off-pathway best fit as reported in Figure 2: k_act_ = 10^8^ M^−1^s^−1^, k_tri_ = 10^6^ M^−1^s^−1^, k_off_ =1.2 s^−1^, N_tri_ = 2, and CF = 1. Note that as k_return_ approaches the magnitude of k_off_, the simulation predicts very little change in the final extent at different pH values, similar to the corresponding linear model.

**Figure S6.**
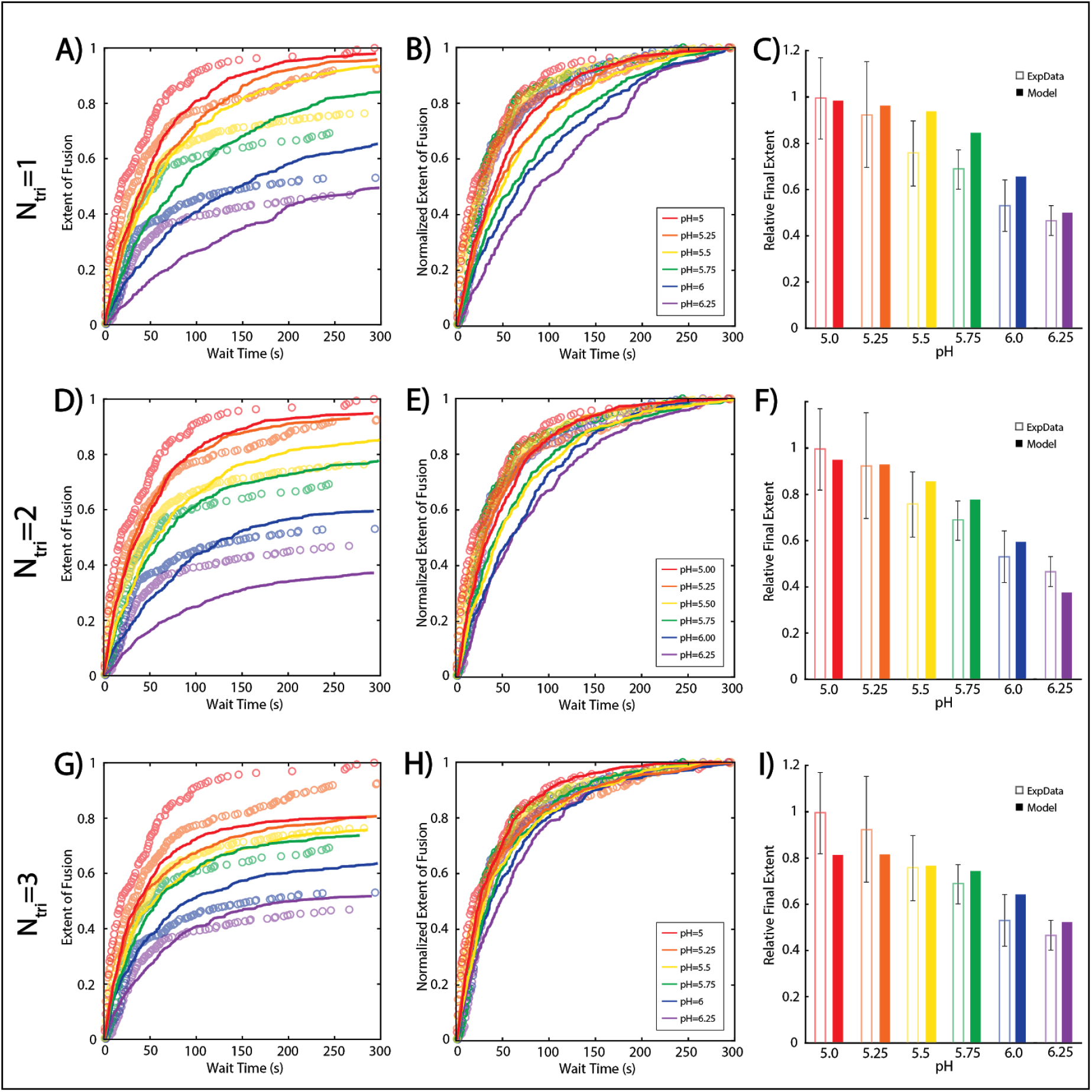
Comparison of CA off-pathway model best fits for N_tri_ = 1, 2, or 3. Single virus hemifusion data was fit to a 5-state off-pathway model of the CA simulation as in Figure 2B. Shown are the best fits for N_tri_ = 1 (Panels A-C) N_tri_ = 2 (Panels D-F), or N_tri_ = 3 (Panels G-I). Best fit parameters for N_tri_ = 1 are: k_act_ = 10^8^ M^−1^s^−1^, k_tri_= 10^6^ M^−1^s^−1^, k_off_= 5.0 s^−1^, f = 0.0047 s^−1^; for N_tri_= 2: k. = 10^8^ M^−1^s^−1^, k_tri_= 10^6^ M^−1^s^−1^, k_off_= 1.2 s^−1^, k_Hfuse_= 0.0047 s^−1^; for N_tri_ = 3: k_act_ = 5 x 10^7^ M^−1^s^−1^, k_tri_= 5 x 10^5^ M^−1^s^−1^, k_off_= 0.25 s^−1^, k_Hfuse_= 0.009 s^−1^.

**Figure S7.**
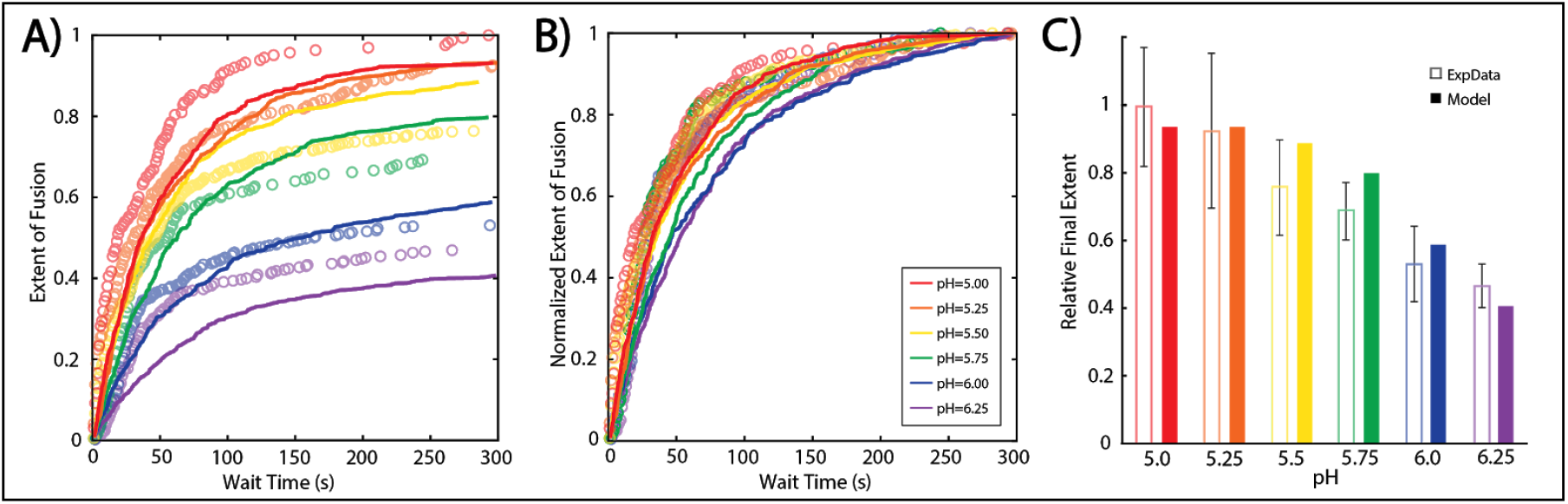
CA off-pathway model best fit at N_tri_ = 2 with varied Cooperativity Factor. Single virus hemifusion data was fit to a 5-state off-pathway model of the CA simulation as in Figure 2B, but varying the Cooperativity Factor in addition to the other free parameters. Best fit parameters with N_trι_=2 are: k_act_ = 10^8^ M^−1^s^−1^, k_trι_= 10^6^ M^−1^s^−1^, k_off_= 1.2 s^−1^, k_Hfuse_= 0.0047 s^−1^, CF=4.

**Figure S8.**
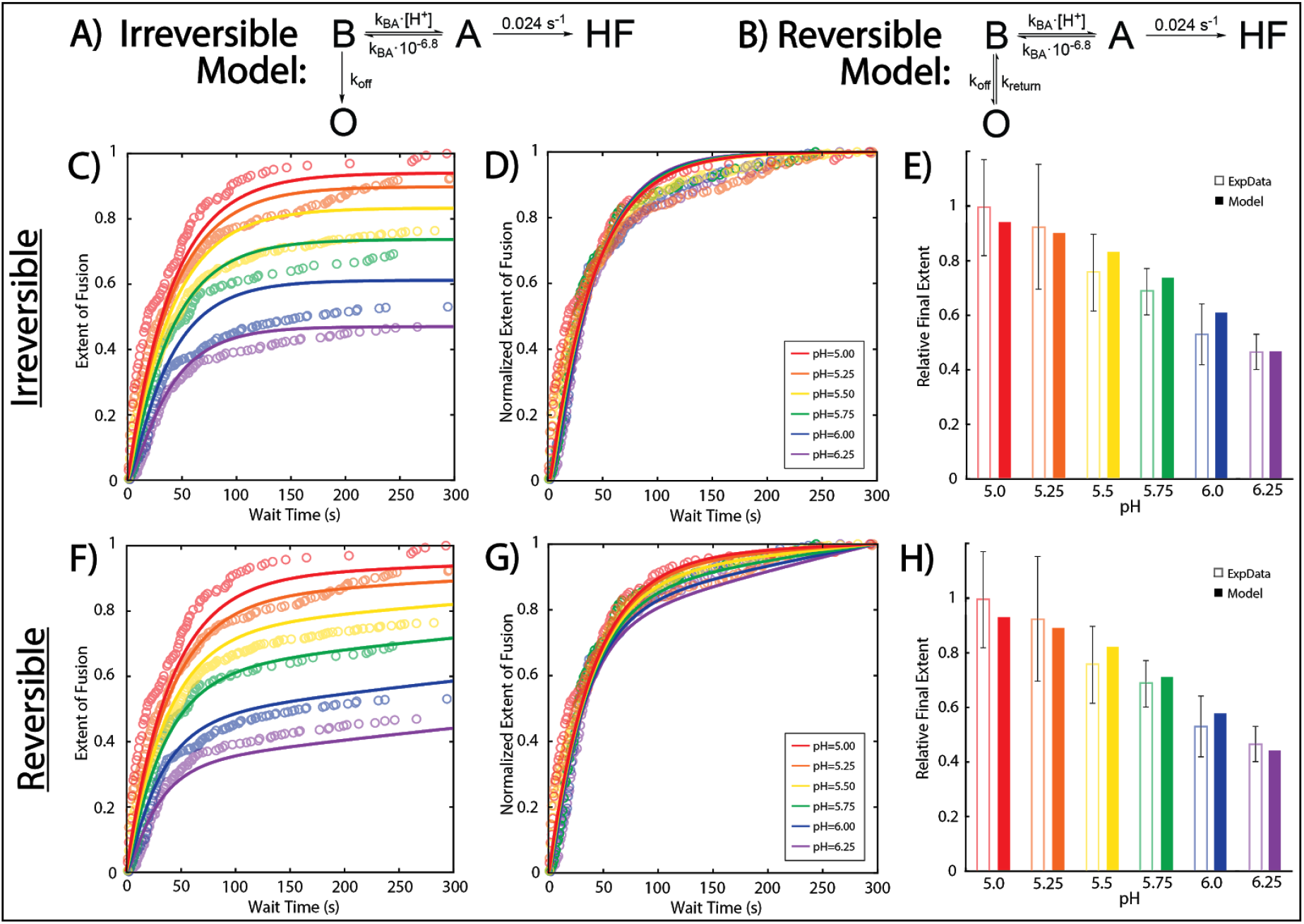
Comparison of irreversible and reversible off-pathway model fits from per virus kinetic modeling. Chemical kinetics-inspired modeling was used to fit the single virus hemifusion data as described in Figure 3, using the off-pathway model with or without k_return_ as a free parameter. The best-fitting parameters for the irreversible model were: k_BA_ = 1.5 x 10^5^ M^−1^s^−1^ and k_off_= 0.05 s^−1^; and for the reversible model were: k_BA_=2.3 x 10^5^ M^−1^s^−1^, k_off_=0.10 s^−1^, and k_return_=1.8 x 10^−4^ s^−1^.

## References

Allison, S.L., Schalich, J., Stiasny, K., Mandl, C.W., Kunz, C., and Heinz, F.X. (1995). Oligomeric rearrangement of tick-borne encephalitis virus envelope proteins induced by an acidic pH. J. Virol. 69, 695–700.

Barrows, N.J., Campos, R.K., Liao, K.-C., Prasanth, K.R., Soto-Acosta, R., Yeh, S.-C., Schott-Lerner, G., Pompon, J., Sessions, O.M., Bradrick, S.S., et al. (2018). Biochemistry and Molecular Biology of Flaviviruses. Chem. Rev. 118, 4448–4482.

Chao, L.H., Klein, D.E., Schmidt, A.G., Peña, J.M., and Harrison, S.C. (2014). Sequential conformational rearrangements in flavivirus membrane fusion. ELife 3, e04389.

Chao, L.H., Jang, J., Johnson, A., Nguyen, A., Gray, N.S., Yang, P.L., and Harrison, S.C. (2018). How small-molecule inhibitors of dengue-virus infection interfere with viral membrane fusion. ELife 7.

Chong, H.Y., Leow, C.Y., Abdul Majeed, A.B., and Leow, C.H. (2019). Flavivirus infection—A review of immunopathogenesis, immunological response, and immunodiagnosis. Virus Res. 274, 197770.

Diamond, M.S. (2009). Virus and Host Determinants of West Nile Virus Pathogenesis. PLoS Pathog. 5, e1000452.

Dowd, K.A., Mukherjee, S., Kuhn, R.J., and Pierson, T.C. (2014). Combined effects of the structural heterogeneity and dynamics of flaviviruses on antibody recognition. J. Virol. 88, 11726–11737.

Floyd, D.L., Ragains, J.R., Skehel, J.J., Harrison, S.C., and van Oijen, A.M. (2008). Single-particle kinetics of influenza virus membrane fusion. Proc. Natl. Acad. Sci. 105, 15382–15387.

Harrison, S.C. (2015). Viral membrane fusion. Virology 479–480, 498–507.

Heinz, F.X., and Stiasny, K. (2012). Flaviviruses and flavivirus vaccines. Vaccine 30, 4301–4306.

Ivanovic, T., and Harrison, S.C. (2015). Distinct functional determinants of influenza hemagglutinin-mediated membrane fusion. ELife 4.

Ivanovic, T., Choi, J.L., Whelan, S.P., van Oijen, A.M., and Harrison, S.C. (2013). Influenza-virus membrane fusion by cooperative fold-back of stochastically induced hemagglutinin intermediates. ELife 2, e00333–e00333.

Kaufmann, B., and Rossmann, M.G. (2011). Molecular mechanisms involved in the early steps of flavivirus cell entry. Microbes Infect. 13, 1–9.

Kim, I.S., Jenni, S., Stanifer, M.L., Roth, E., Whelan, S.P.J., van Oijen, A.M., and Harrison, S.C. (2017). Mechanism of membrane fusion induced by vesicular stomatitis virus G protein. Proc. Natl. Acad. Sci. 114, E28–E36.

Krishnan, M.N., Sukumaran, B., Pal, U., Agaisse, H., Murray, J.L., Hodge, T.W., and Fikrig, E. (2007). Rab 5 is required for the cellular entry of dengue and West Nile viruses. J. Virol. 81, 4881–4885.

Kuhn, R.J., Dowd, K.A., Beth Post, C., and Pierson, T.C. (2015). Shake, rattle, and roll: Impact of the dynamics of flavivirus particles on their interactions with the host. Virology 479–480, 508–517.

Liu, K.N., and Boxer, S.G. (2020). Target Membrane Cholesterol Modulates Single Influenza Virus Membrane Fusion Efficiency but Not Rate. Biophys. J. S000634952030271X.

Otterstrom, J.J., Brandenburg, B., Koldijk, M.H., Juraszek, J., Tang, C., Mashaghi, S., Kwaks, T., Goudsmit, J., Vogels, R., Friesen, R.H.E., et al. (2014). Relating influenza virus membrane fusion kinetics to stoichiometry of neutralizing antibodies at the single-particle level. Proc. Natl. Acad. Sci. 111, E5143–E5148.

Perera-Lecoin, M., Meertens, L., Carnec, X., and Amara, A. (2013). Flavivirus Entry Receptors: An Update. Viruses 6, 69–88.

Rawle, R.J., Boxer, S.G., and Kasson, P.M. (2016). Disentangling Viral Membrane Fusion from Receptor Binding Using Synthetic DNA-Lipid Conjugates. Biophys. J. 111, 123–131.

Rawle, R.J., Webster, E.R., Jelen, M., Kasson, P.M., and Boxer, S.G. (2018). pH Dependence of Zika Membrane Fusion Kinetics Reveals an Off-Pathway State. ACS Cent. Sci. 4, 1503–1510.

Rawle, R.J., Villamil Giraldo, A.M., Boxer, S.G., and Kasson, P.M. (2019). Detecting and Controlling Dye Effects in Single-Virus Fusion Experiments. Biophys. J.

van der Schaar, H.M., Rust, M.J., Waarts, B.-L., van der Ende-Metselaar, H., Kuhn, R.J., Wilschut, J., Zhuang, X., and Smit, J.M. (2007). Characterization of the Early Events in Dengue Virus Cell Entry by Biochemical Assays and Single-Virus Tracking. J. Virol. 81, 12019–12028.

Schmidt, A.G., Lee, K., Yang, P.L., and Harrison, S.C. (2012). Small-molecule inhibitors of dengue-virus entry. PLoS Pathog. 8, e1002627.

Stiasny, K., and Heinz, F.X. (2004). Effect of Membrane Curvature-Modifying Lipids on Membrane Fusion by Tick-Borne Encephalitis Virus. J. Virol. 78, 8536–8542.

de Wispelaere, M., Lian, W., Potisopon, S., Li, P.-C., Jang, J., Ficarro, S.B., Clark, M.J., Zhu, X., Kaplan, J.B., Pitts, J.D., et al. (2018). Inhibition of Flaviviruses by Targeting a Conserved Pocket on the Viral Envelope Protein. Cell Chem. Biol. 25, 1006–1016.e8.

Yang, S.-T., Kiessling, V., Simmons, J.A., White, J.M., and Tamm, L.K. (2015). HIV gp41–mediated membrane fusion occurs at edges of cholesterol-rich lipid domains. Nat. Chem. Biol.

Yang, S.-T., Kreutzberger, A.J.B., Kiessling, V., Ganser-Pornillos, B.K., White, J.M., and Tamm, L.K. (2017). HIV virions sense plasma membrane heterogeneity for cell entry. Sci. Adv. 3, e1700338.

